# SEC62 and TMX4 control asymmetric autophagy of the nuclear envelope upon LINC complex disassembly

**DOI:** 10.1101/2022.10.05.510950

**Authors:** Marika K. Kucińska, Juliette Fedry, Carmela Galli, Diego Morone, Andrea Raimondi, Tatiana Soldà, Friedrich Förster, Maurizio Molinari

## Abstract

The endoplasmic reticulum (ER) is a dynamic organelle of nucleated cells that produces proteins, lipids and oligosaccharides. The volume and the activities of the ER are adapted to cellular needs. They are increased upon induction of unfolded protein responses (UPR ^1^) and are reduced upon activation of ER-phagy programs ^2^. A specialized domain of the ER, the nuclear envelope (NE), protects the cell genome with two juxtaposed lipid bilayers, the inner and outer nuclear membranes (INM and ONM). SUN proteins in the INM form disulfide-bonded Linker of Nucleoskeleton and Cytoskeleton (LINC) complexes with NESPRIN proteins in the ONM. These complexes set and maintain the width of the periplasmic space (PS), a continuum of the ER lumen, below the 50 nm ^3-5^ and in yeast prevent transmission of ER volume variations to the PS ^6,7^. Here we report that expansion of the mammalian ER upon homeostatic perturbations is transmitted to the NE, where the ONM forms large bulges. The process is reverted on recovery of ER homeostasis, by asymmetric vesiculation and autophagic clearance of ONM portions. Remodeling of the mammalian NE requires TMX4-driven reduction of the *inter*molecular disulfide bond stabilizing LINC complexes, the LC3 lipidation machinery, and the autophagy receptor SEC62, identified here as the first mammalian nucleo-phagy receptor.

---

To assess a possible transmission of the ER swelling elicited by a pharmacologic treatment that perturbs ER homeostasis to the NE, we exposed mouse embryonic fibroblasts (MEF), NIH/3T3 MEF, and human embryonic kidney (HEK293) cells, to cyclopiazonic acid (CPA), a reversible inhibitor of the sarco/ER calcium pump ^8-10^. Low CPA dosages mimic ER stresses triggered by perturbation of glycosylation (tunicamycin ^11,12^), redox (DTT ^9,12,13^), or calcium homeostasis (thapsigargin ^12,14^ or phenobarbital ^15-17^), which are characterized by ER expansion as well as transcriptional and translational induction of ER-resident proteins ^8-10^. The advantage of CPA administration is that the compound is non-toxic at low dosage, and the ER stress is fully reversible upon interruption of the pharmacologic treatment ^8-10^.

Dynamic changes in NE shape have been monitored by cryogenic Focus Ion Beam milling followed by Cryo Electron Tomography (FIB-CET ^18^) on vitrified cells (**Figs. 1B-1G**). Analyses of 17 tomograms reveals that at steady state, the distance between ONM and INM generally remains below 50 nm (**Fig. 1D** and **SFig. 1A**, pale blue zone). This distribution is expected because the distance between INM and ONM, i.e., the width of the PS, is determined by the maximal extension of the elastic perinuclear domains of SUN proteins within LINC complexes (**Fig. 1A** and Inset) ^3-5,19,20^. Exposure of mammalian cells to 10 µM CPA for 12 h substantially increases the median distance between the INM and the ONM (**Figs. 1E-1G** and **SFig. 1B**). The increased median distance between INM and ONM upon induction of a mild ER stress was also observed by morphometric analyses of the NE by Room Temperature-Transmission Electron Microscopy (RT-TEM) of chemically fixed, resin embedded cells (**Figs. 1H** *vs*. **1I** and **SFigs. 2A** *vs*. **2B**). Inspection of the images reveals that the ONM swells and forms bulges upon perturbation of ER homeostasis (**SFig. 2B**). These results highlight a major difference with yeast cells (*Saccharomyces cerevisiae*), where ER expansion is not transmitted to the NE, and LINC complexes stabilize the PS volume ^3-7^.

**Fig. 1.**
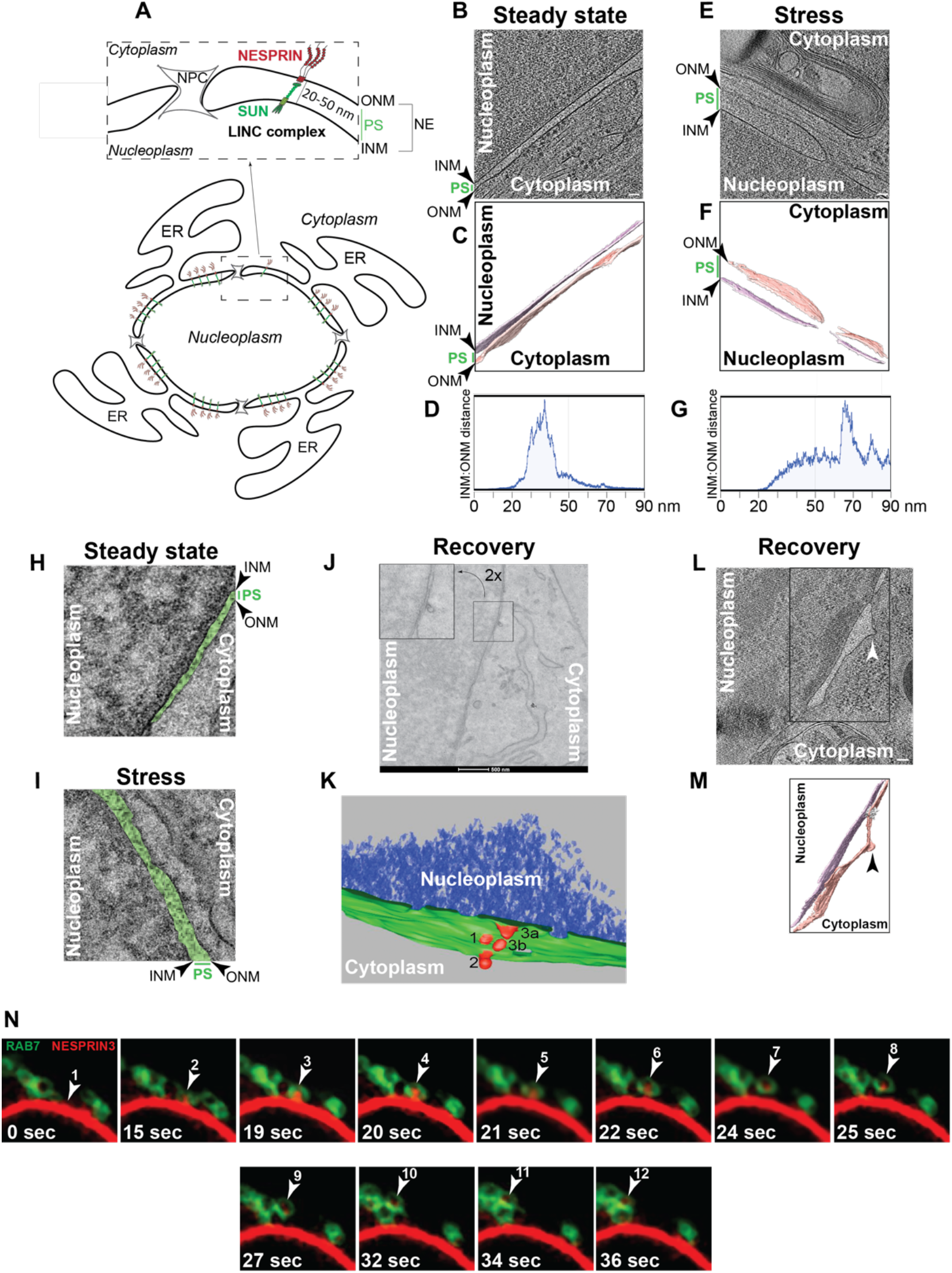
NE dynamics and vesiculation. **A** Schematic representation of the NE, with inset to show the LINC complex, which maintains the distance between ONM and INM below the 50 nm. **B** Slice through example CET of NEs from a lamella through a cell at steady state. Scale bar, 50 nm. **C** Isosurface representation of the corresponding segmented volumes for NE at steady state. ONM in salmon; INM in light purple. **D** Quantifications of corresponding ONM-INM distances at steady state (see Tomogram 14, **SFig1A**). **E** Same as **B** under ER stress. Scale bar, 50 nm. **F** Same as **C** under ER stress. **G** Same as **D** for a cell under ER stress (see Tomogram 1, **SFig. 1B**). **H** TEM micrograph of a portion of NE in MEF at steady state (the periplasmic space (PS) is in green). **I** Same as **H** for MEF under ER stress. **J** Same as **H** and **I** for MEF after 5 h recovery. **K** RT-ET of a subdomain of the NE in a recovering MEF reconstructed by TEM Tomography shows an initial budding of the ONM (1), a later stage of the process of vesicle formation (2) and the final step where the vesicle has just been released from the ONM (3) occur in proximity. Please also refer to **Movie 1**. **L** Slice through a CET of the NE of a recovering MEF. Scale bar corresponds to 100 nm. White arrow indicates the formation of a budding site at the ONM. Please also refer to **Movie 2**. **M** Isosurface representation of the corresponding segmented volume. ONM in salmon; INM in light purple. **N** Frames of a movie (**Movie 3**) showing the capture of a NESPRIN3-positive ONM-derived vesicle by a GFP-RAB7-positive endolysosome.

**Fig. 2.**
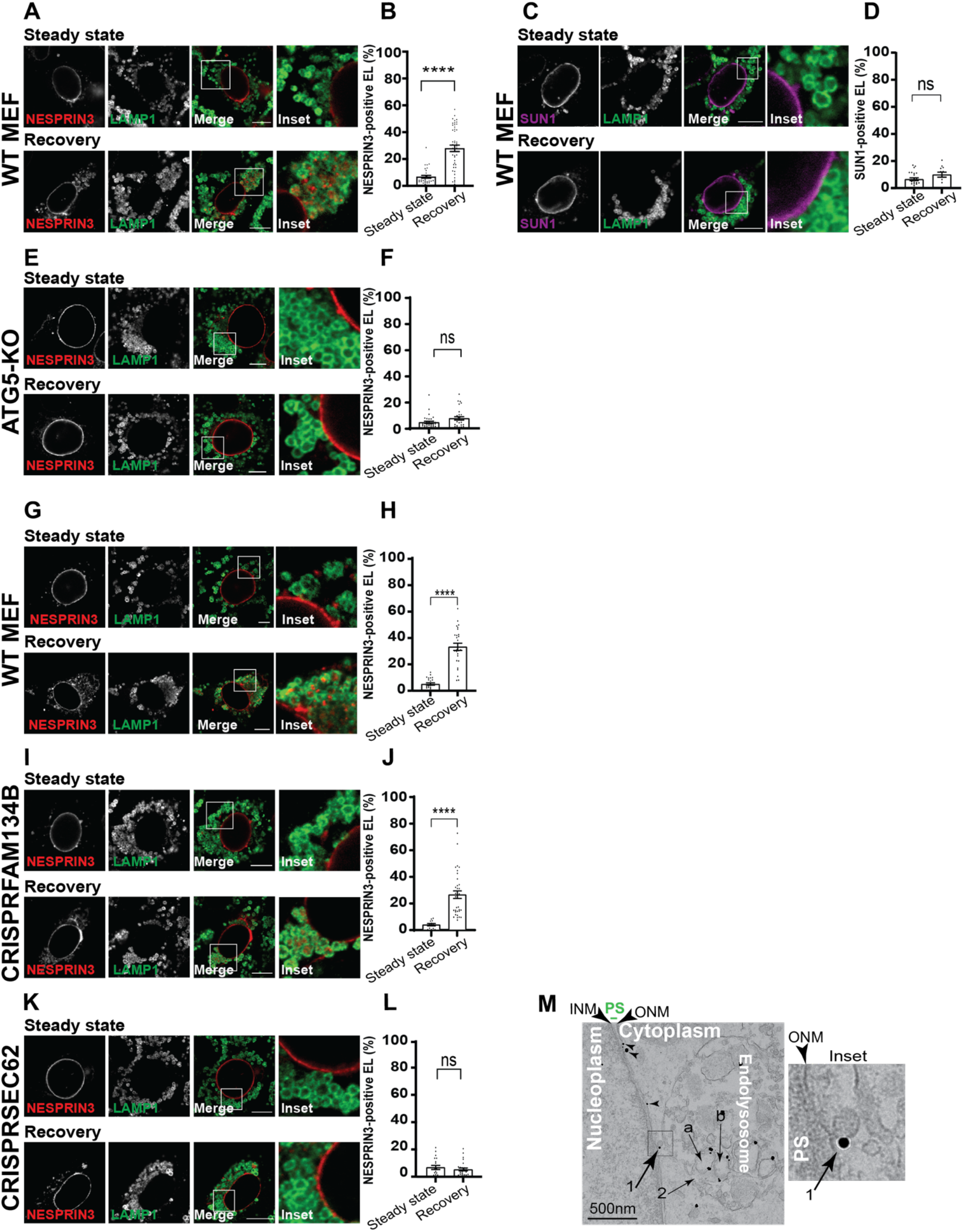
Lysosomal delivery of ONM subdomains. **A** CLSM showing the sub-cellular localization of NESPRIN3 in WT MEF at steady state (upper panels) and 12 h after interruption of the pharmacologic treatment (lower panels). Endolysosomes are stained with a LAMP1 antibody. Note that cells were exposed to the lysosomal inhibitor bafilomycinA1 (BafA1) to prevent clearance of the INM and ONM marker proteins if delivered within endolysosomes. **B** Quantification of **A**. n =3 independent experiments. mean ± SEM; unpaired, two-tailed t test, ****P< 0.0001 **C** Same as **A**, to show the sub-cellular localization of SUN1. **D** Quantification of **C**. n = 3 independent experiments, mean ± SEM; unpaired, two-tailed t test, ns. P= 0.1007 **E** Same as **A** in cells lacking the autophagy gene product ATG5 ^9^. **F** Quantification of **E**. n = 2 independent experiments, mean ± SEM; unpaired, two-tailed t test, ns. P=0.0543 **G** Same as **A** in WT MEF. **H** Quantification of **G**. n = 3 independent experiments, mean ± SEM; unpaired, two-tailed t test, ***P< <0.0001 **I** Same as **G** in cells lacking the ER-phagy receptor FAM134B ^9,10^. **J** Quantification of **I**. n = 2 independent experiments, mean ± SEM; unpaired, two-tailed t test, ****P< 0.0001 **K** Same as **G** in cells lacking the autophagy receptor SEC62 ^9,10^. **L** Quantification of **K**. n = 3 independent experiments, mean ± SEM; unpaired, two-tailed t test, ns. P=0.4279. Scale bars, 10 μm. **M** Immunogold EM to label endogenous SEC62 in MEF recovering from ER stress. Budding of a SEC62-positive vesicles from the ONM (arrow 1). Endolysosome capturing SEC62-labeled vesicles (arrow 2).

As previously observed for other medical compounds whose administration causes ER swelling, most notably the anti-epileptic drug phenobarbital ^15-17^, interruption of the pharmacologic treatment with CPA activates autophagy programs relying on vesiculation of ER subdomains and their delivery to endolysosomal compartments for clearance ^9,10^, to resume the pre-stress ER volume and activity ^9,10,15,16^. We reasoned that conclusion of the ER stress could also trigger autophagic programs to re-establish the pre-stress distance between INM and ONM, thereby returning the periplasmic volume to physiologic levels.

To verify this hypothesis, CPA-treatment was interrupted to initiate the recovery phase. Five hours later, cells were processed for imaging. RT-TEM micrographs (**Figs. 1J, SFig. 2C**, Insets) Room Temperature-Electron Tomography (RT-ET) (**Fig. 1K, Movie 1**) and CET volumes (**Figs. 1L, 1M**, arrow, **Movie 2**) reveal that during the recovery phase, portions of the ONM project into the cytoplasm and vesiculate. Regions of the NE are seen in RT-ET, where the initial phase of ONM budding (**Fig. 1K**, 1 and **Movie 1**), the closure of the bud that precedes detachment of the ONM-derived vesicle (**Fig. 1K**, 2 and **Movie 1**), and the actual detachment of a vesicle (**Fig. 1K**, 3a and 3b, **Movie 1**), are simultaneously proceeding in a delimited domain of the ONM. This series of events is dissected in time-course analyses by confocal light scanning microscopy (CLSM) of living cells (selected frames in **Fig. 1N** and **Movie 3**). Live cell imaging confirms the formation of buds and vesicles containing NESPRIN3, a protein marker of the ONM ^21^ (**SFig. 3A**), which are engulfed by GFP-RAB7-positive endolysosomes for clearance (**Fig. 1N, Movie 3**).

**Fig. 3.**
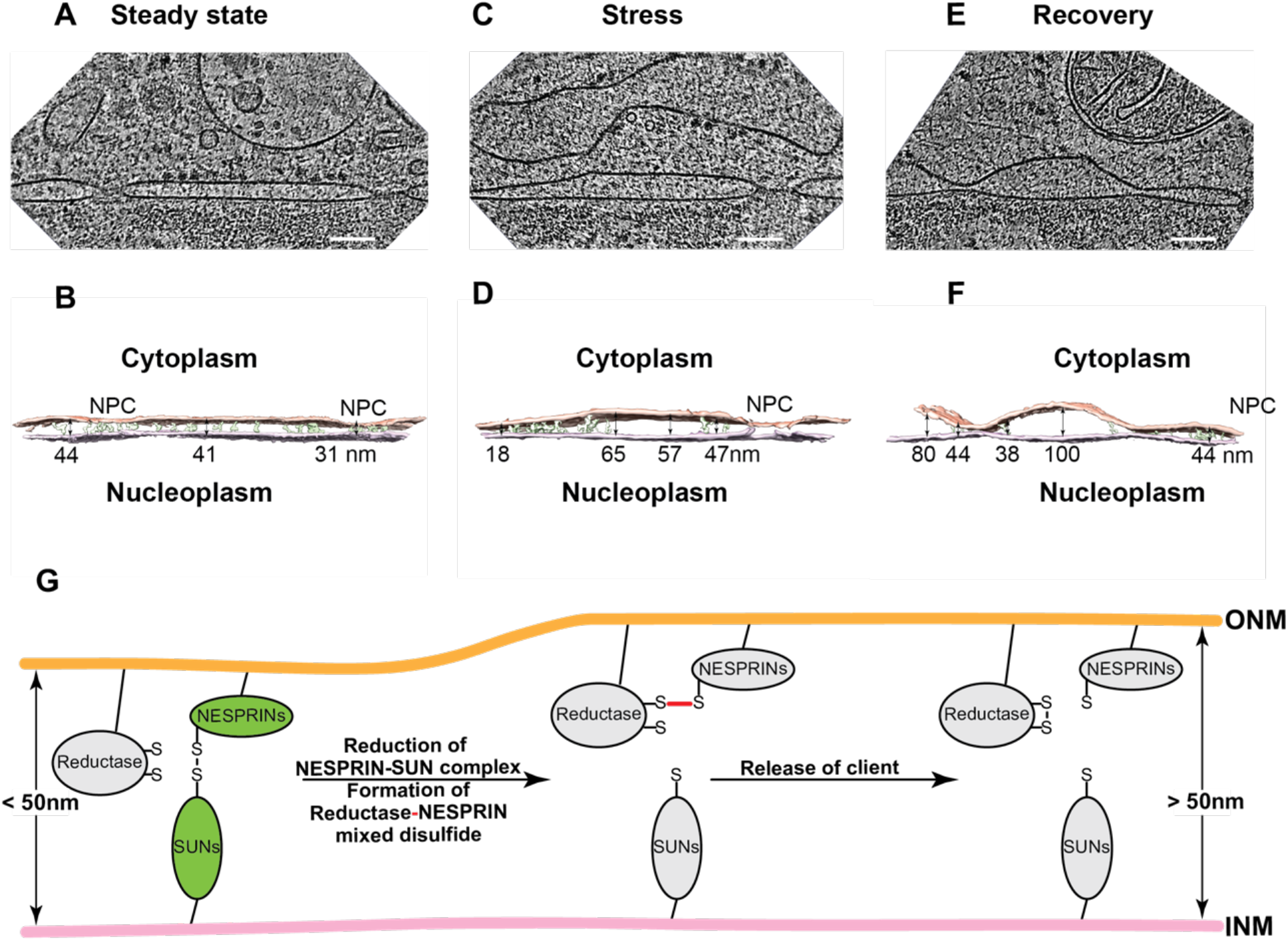
Visualizing LINC complex filaments. **A** Slice through cryo electron tomogram of a NE in a FIB-milled MEF at Steady state. Scale bar represents 100 nm. **B** Isosurface representation of the corresponding segmented volume (ONM in salmon, INM in light purple). Filaments traced in the perinuclear space are depicted in green (continuous from ONM to INM). Indicative distances between ONM and INM are given along the membrane. **C** Same as **A** for cells under stress. **D** Same as **B**. **E** Same as **A** for recovering cells. **F** Same as **B**. **G** Proposed mode-of-action of a reductase that engages NESPRINs in mixed disulfides (red bond) to eventually disassemble the LINC complex.

To further demonstrate the asymmetric vesiculation of the NE and the selective delivery of ONM portions to endolysosomal compartments for clearance during recovery from ER stress (**Fig. 1**), we have monitored in CLSM the intracellular fate of HALO-tagged NESPRIN3 and of GFP-tagged SUN1 selected as protein markers of the ONM and of the INM, respectively (**SFig. 3A**) ^21^. At steady state, both proteins are in the NE (**Figs. 2A** and **2C**, upper panels, **SFig. 3A**) ^21^ and are excluded from LAMP1-positive endolysosomes (**Figs. 2A** and **2C**, upper panels, Insets). Thus, at steady state the delivery of these ONM and INM protein markers within degradative compartments is below detection level. During the recovery phase initiated upon interruption of the pharmacologic treatment with CPA, a substantial fraction of NESPRIN3 re-localizes in cytoplasmic puncta (**Fig. 2A**, lower panel, NESPRIN3), within LAMP1-positive endolysosomes, where it accumulates upon inactivation of the lysosomal hydrolytic activity (**Fig. 2A**, lower panels, Inset). Instead, the INM marker protein SUN1 is not delivered to the lysosomal compartment (**Fig. 2C**, lower panels, Inset) as quantified using LysoQuant, an unbiased and automated deep learning image analysis tool for segmentation and classification of fluorescence images capturing cargo delivery to endolysosomes ^22^ (**Figs. 2B** and **2D**). Delivery of NESPRIN3, but not of SUN1, to the endolysosomal compartment for clearance confirms the asymmetric autophagy of the ONM during recovery from ER stress (**Figs. 1J-1N, SFig. 2C, Movie 3**).

Vesiculation and lysosomal delivery of organelle portions for clearance involves autophagy gene products, most notably those regulating lipidation of the ubiquitin-like protein LC3, and organelle-specific LC3-binding autophagy receptors displayed at the limiting membrane of the organelle portion to be removed from cells ^2,23,24^. Inspired by these findings, we first assessed lysosomal delivery of NESPRIN3 in cells lacking ATG5, i.e., defective in LC3 lipidation ^25^. Like WT MEF (**Figs. 2A, 2B**), the absence of NESPRIN3 immunoreactivity in LAMP1-positive endolysosomes confirms lack of detectable constitutive delivery of ONM to degradative compartments at steady state (**Figs. 2E**, upper panels, **2F**, Steady state). As reported for other types of organello-phagy ^2,23,24^, LysoQuant analyses reveal that the absence of ATG5 substantially inhibits delivery of NESPRIN3-positive ONM portions to the lysosomal compartments, which should be activated during the recovery phase from ER stress (**Figs. 2E**, lower panels, **2F**, Recovery).

In yeast, selective autophagy of nuclear components (i.e., nucleus-derived double membrane vesicles, NE domains containing specific nucleoporins, nuclear pore complexes) has mainly been studied as a catabolic process induced upon nutrient deprivation and it involves the autophagy receptor Atg39 ^26-29^. In mammalian cells, selective autophagy of nuclear components has been observed in response to oncogenic insults ^30^, DNA damage ^31^ and cellular senescence ^32^. However, the molecular mechanisms of these catabolic processes has not been elucidated, and a functional ortholog of Atg39 that regulates autophagy of nuclear portions remains to be characterized ^27^. To identify the autophagy receptor controlling delivery of ONM-derived vesicles to the endolysosomes for clearance, we reasoned that recovery from acute ER stresses activates the function of SEC62 as autophagy receptor ^9,10^. Relevantly, SEC62 localizes in the ONM (**Fig. 2M**, arrowheads, **SFigs. 3B, 3C**).

To assess the involvement of SEC62 as the autophagy receptor regulating lysosomal clearance of the NESPRIN3-positive ONM portions, we compared lysosomal delivery of NESPRIN3 to the LAMP1-positive endolysosomes in WT MEF, in MEF lacking the starvation-induced autophagy receptor FAM134B, and in cells lacking SEC62 generated by CRISPR-Cas9 genome editing ^9,10^. The CLSM and LysoQuant analyses confirm the increased delivery of NESPRIN3 to the LAMP1-positive degradative compartment in WT MEF recovering from ER stress (**Figs. 2G, 2H**, Steady state *vs*. Recovery). Deletion of the ER-phagy receptor FAM134B does not affect delivery of NESPRIN3 to the endolysosomal compartment during recovery from ER stress showing dispensability of this autophagy receptor for lysosomal turnover of ONM subdomains (**Figs. 2I, 2J**). In contrast, deletion of SEC62 substantially inhibits the activation of NESPRIN3 delivery to the lysosomal compartment during Recovery (**Figs. 2K, 2L**, Steady state *vs*. Recovery).

Notably, SEC62 is displayed at the limiting membrane of ONM-derived vesicles (**Fig. 2M** and Inset, arrow 1), and these vesicles may be seen in proximity to endolysosomes (**Fig. 2M**), which eventually capture them for clearance as shown in live cell imaging (**Fig. 1N**). Our data show that the autophagy receptor SEC62, which is activated during recovery from ER stress to regulate return of ER volume and content at physiologic levels ^9,10^, is also involved in selection of ONM subdomains for lysosomal clearance.

Analyses of the micrographs to compare the morphology of the NE in cells at steady state (**SFig. 2A**), in cells with pharmacologically perturbed ER homeostasis (**SFig. 2B**) and in cells restoring physiologic conditions (**SFig. 2C**), show that the ONM is more irregular in the two latter cases. In these cells, bulges are formed, where the distance from the INM can reach the 120 nm and more (**Fig. 1G, SFigs. 1B, 1C**). The integrity of the NE is ensured by the LINC complexes, which are composed of INM proteins of the SUN family and ONM proteins of the NESPRIN family (**Fig. 1A**). SUNs and NESPRINs are covalently linked, head-to-tail, by *inter*molecular disulfide-bonds. Visualization of native NE of MEF at steady state (**Figs. 3A, 3B**) exposed to ER stress (**Figs. 3C, 3D**) or during recovery from ER stress (**Figs. 3E, 3F**) by FIB-CET and manual density-based segmentation reveals continuous periplasmic densities from ONM to INM only in NE subdomains, where the distance between the lipid bilayers is below the 55 nm (**Figs. 3B, 3D, 3F**). Above this limit, filaments connecting the INM and the ONM are not visible. This observation is in good agreement with structural data on the elastic perinuclear domains of SUN trimers, which determine the distance between the two lipid bilayers of the NE, maximally elongating up to 50 nm ^3-5^. The frequent occurrence of bulges with distances above the 50 nm during the ER stress and the recovery phases (**SFigs. 1B, 1C**), and the asymmetric vesiculation of the ONM during recovery from the ER stress (**SFig. 1C**) imply an enzyme-catalyzed pathway that disassembles LINC complexes upon reduction of the structural, *inter*molecular disulfide bond linking the periplasmic domains of SUN and NESPRIN proteins (**Fig. 3G**).

More than 20 oxidoreductases of the protein disulfide isomerase (PDI) superfamily populate the ER ^33^ (endogenous PDI, ERp57, ERp72 and ectopically expressed TMX3-V5 are shown as examples, **SFigs. 3D-3G**, respectively). One of the oxidoreductases we tested, TMX4, has a peculiar enrichment in the NE (**Fig. 4A**) that has been reported before ^34^. TMX4 has a non-canonical cysteine-proline-serine-cysteine (*CPSC)* active site sequence. The proline residue at position 2 destabilizes the disulfide state and favors the di-thiol reduced form of the active site ^35^. Consistently, TMX4 acts as reductase *in vitro* ^36^, but its biological function remains to be established ^37,38^. The peculiar sub-cellular distribution of TMX4 in the NE, and its reported *in vitro* activity as a reductase, led us to check if TMX4 interacts with NESPRIN3 and regulates the redox-driven disassembly of LINC complexes. We first assessed whether TMX4 engages NESPRIN3 in mixed disulfides, an obligate reaction intermediate during disulfide-bonded complex disassembly (**Fig. 3C**, red bond) ^33,39^. To this end, HALO-tagged NESPRIN3 was expressed in MEF alone (**Fig. 4B**, lane 1), or with V5-tagged TMX4_C67A_ *trapping mutant* (**Fig. 4B**, lane 3). The replacement of the second cysteine residue in the _64_CPSC_67_ catalytic site of TMX4 with an alanine residue stabilizes mixed disulfide species preventing release of TMX4 clients ^40^.

**Fig. 4.**
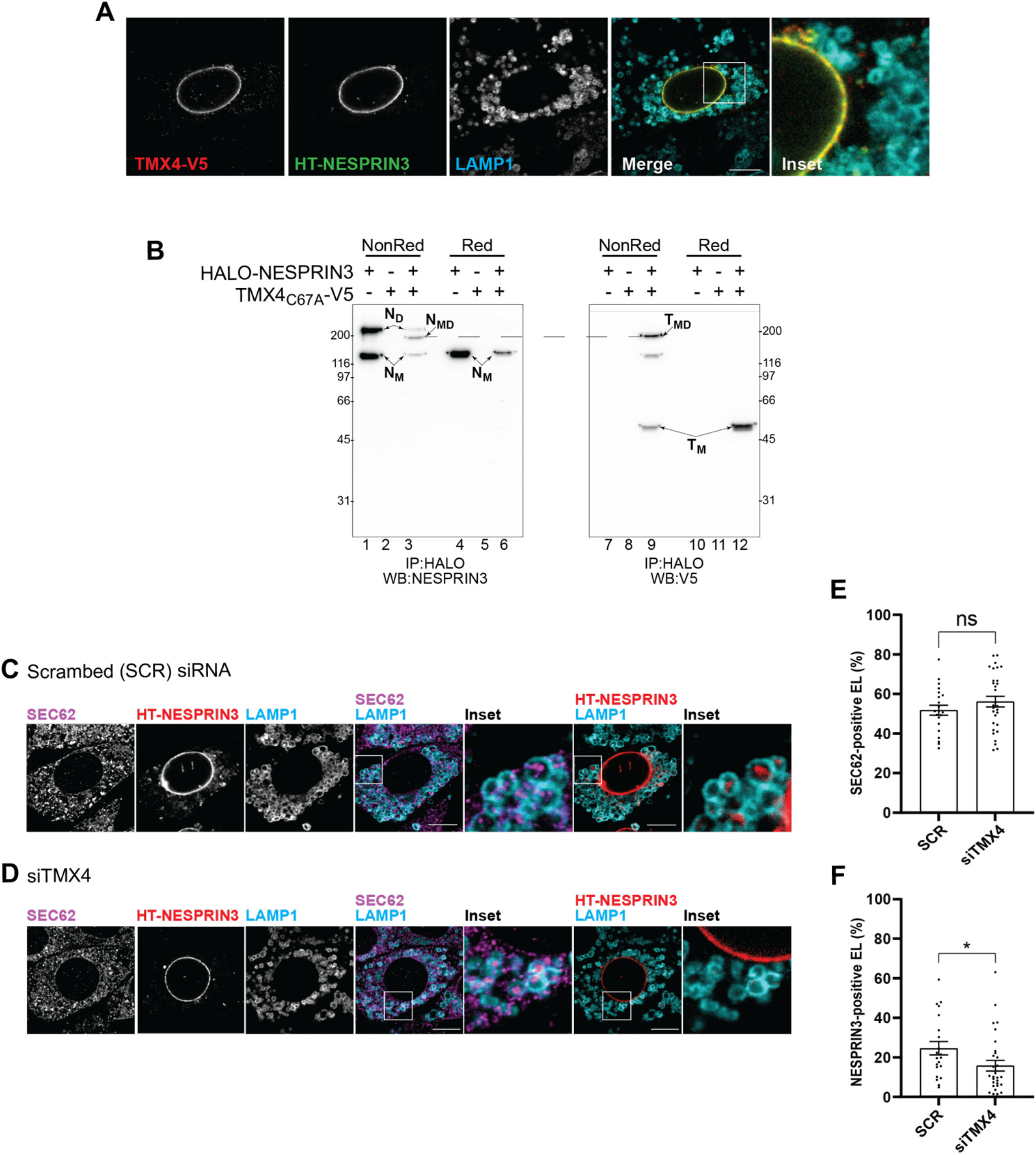
Involvement of TMX4 in ONM-phagy. **A** Intracellular expression of TMX4-V5, which co-localizes with HALO-NESPRIN3 in the NE. **B** Analyses of disulfide-bonded complexes formed *in cellula* when HALO-NESPRIN3 is expressed alone (lanes 1, 4, 7, 10), or is co-expressed with TMX4_C67A_-V5 (lanes 3, 6, 9, 12). In lanes 2, 5, 8, 11, cells expressing only TMX4_C67A_-V5. In lanes 1-3 and 7-9 the polypeptides were separated under non reducing conditions to preserve disulfide-bonded complexes. In lanes 4-6 and 10-12 they were separated under reducing conditions to disassemble the complexes. Lanes 1-6 show the NESPRIN3 component of the complexes (N_M_, N_D_, N_MD_ for NESPRIN3 Monomers, Dimers and Mixed Disulfides (engaging TMX4), respectively). Lanes 7-12 show the TMX4 component of the complexes (T_M_ and T_MD_ for TMX4 monomers and Mixed Disulfides (engaging NESPRIN3), respectively). **C** Same as **Fig. 2**. CLSM showing the delivery of endogenous SEC62 and HALO-NESPRIN3 within LAMP1-positive endolysosomes during recovery from stress. **D** Same as H, in cells where TMX4 expression has been silenced by RNA interference. **E** Quantification of SEC62-positive endolysosomes in mock-treated cells and in cells with reduced expression of TMX4. **F** Same as J for NESPRIN3-positive endolysosomes. Scale bars, 10 μm.

Ectopically expressed NESPRIN3 was immunoisolated from cell lysates with anti-HALO antibodies and separated in non-reducing (**Fig. 4B**, lanes 1-3), or in reducing SDS-polyacrylamide gels (lanes 4-6). The gels were processed for western blotting with anti-NESPRIN3 antibodies to revel the NESPRIN3 component of mixed disulfides (**Fig. 4B**, lanes 1-6), or with anti-V5 antibodies to reveal the TMX4 component of mixed disulfides (lanes 7-12). Under non-reducing conditions, HALO-NESPRIN3 is separated into two polypeptide bands of approximately 140 and 280 kDa, respectively (**Fig. 4B**, lane 1, arrows N_M_ for <u>N</u>ESPRIN3 <u>M</u>onomers and N_D_ for <u>N</u>ESPRIN3 <u>D</u>imers). In reducing gels, monomers and disulfide-bonded dimers of HALO-NESPRIN3 migrate as a single polypeptide upon reduction of the *inter*molecular disulfide bonds (**Fig. 4B**, lane 4, arrow N_M_). Upon co-expression with TMX4_C67A_-V5 (lane 3), an additional polypeptide band at approximately 200 kDa shows up (**Fig. 4B**, lane 3, N_MD_). This band corresponds to TMX4-S-S-NESPRIN3 mixed disulfide as proven by the immunoreactivity with anti-NESPRIN3 antibodies that reveal the <u>N</u>ESPRIN3 component of <u>M</u>ixed <u>D</u>isulfides (N_MD_ in lane 3), and with V5-specific antibodies that reveals the <u>T</u>MX4 component of <u>M</u>ixed <u>D</u>isulfides (**Fig. 4B**, T_MD_ in lane 9). As expected, the two components are dissociated under reducing conditions, where the signal collapses in the NESPRIN3 (**Fig. 4B**, lane 6, N_M_) and TMX4-immunoreactive polypeptides (**Fig. 4B**, lane 12, T_M_).

Next, we assessed whether TMX4 engages endogenous NESPRIN proteins in mixed disulfides *in cellula*. To this end, V5-tagged TMX4, or its trapping mutant version TMX4_C67A_ were transiently expressed in MEF. As specificity control, MEF were transfected with plasmids for expression of TMX3 or of the corresponding TMX3_C56A_ trapping mutant. TMX3 is a TMX4 paralog, prevalently located in the ER ^41^ (**SFig. 3G**), where it catalyzes oxidation of client proteins ^41^. After immunoisolation from cell lysates with anti-V5 antibodies, the immunocomplexes containing TMX3-V5 (**SFig. 4A**, lanes 2 and 8), TMX3_C56A_-V5 (lanes 3 and 9), TMX4-V5 (lanes 4 and 10), TMX4_C67A_-V5 (lanes 5 and 11) were separated in non-reducing/reducing SDS-polyacrylamide gel, which was subsequently silver stained (**SFig. 4A**). TMX3_C56A_ and TMX4_C67A_ are trapped in mixed disulfides with several cellular polypeptides (**SFig. 4A**, lanes 3 and 5, respectively, red rectangles). These disulfide-bonded complexes disappear from the region of the gels shown with the red rectangles upon reduction (**SFig. 4A**, lanes 9 and 11). As expected, the disulfide-bonded complexes are significantly less abundant in the lanes loaded with immunoisolates of cells expressing the active forms of TMX3 and TMX4, which rapidly release their clients (**SFig. 4A**, lanes 2 and 4, black rectangles).

The cellular proteins captured in disulfide bonded complexes with the TMX3 and the TMX4 trapping mutants (**SFig. 4A**, lanes 3 and 5, respectively, red rectangles) were identified by mass spectrometry. These analyses reveal endogenous NESPRIN proteins (i.e., NESPRIN2 and NESPRIN1) as major TMX4 clients (**SFig. 4B**, blue). NESPRIN proteins were not captured by the TMX3_C56A_ trapping mutant showing a different client subset for the two PDI family members. A previously published study performed to identify clients of TMX1 ^42^, and unpublished data from our group aiming at the identification of the clients of TMX5 (**SFig. 4A**, lanes 6 and 12), two other membrane-bound members of the PDI superfamily, also failed to detect NESPRINs amongst TMX1 ^42^ or TMX5 clients (unpublished). Thus, TMX4 has peculiar localization among PDI family members in the NE (**Fig. 4A**) ^34^, has reductase activity ^36^, preferentially associates with endogenous NESPRIN proteins in living cells (**SFig. 4**), with whom it can be trapped in mixed disulfides (**Fig. 4B**) as an indication of an involvement of TMX4 as regulator of redox cycles determining NESPRIN3 engagement in supramolecular complexes.

Last, during recovery from acute ER stresses, ER subdomains ^9,10^ and ONM portions (this work) displaying the autophagy receptor SEC62 at the limiting membrane are delivered to LAMP1-positive endolysosomes for clearance. This is once more recapitulated in MEF cells transfected with scrambled small interfering (si)RNA (SCR), where SEC62 and NESPRIN3 are delivered within LAMP1-positive endolysosomes during recovery from ER stress (**Figs. 4C** and Insets, **4E** and **4F**, SCR). Silencing of TMX4 expression by about 70% (siTMX4, **SFigs. 3H, 3I**) does not affect lysosomal delivery of SEC62 (**Figs. 4D**, SEC62/LAMP1 and Inset, **4E**, SCR *vs*. siTMX4), but inhibits lysosomal delivery of NESPRIN3 (**Figs. 4D**, NESPRIN3/LAMP1 and Inset, **Fig. 4F**, SCR *vs*. siTMX4). This observation hints at a mechanistic difference between lysosomal clearance of ER subdomains ^9,10^ and of ONM portions. Both pathways involve the autophagy receptor SEC62, but only the lysosomal delivery of ONM requires a TMX4-driven redox event that segregates ONM portions to be removed from cells from the INM, which is preserved, possibly to ensure better protection of the genetic material with which it is in direct contact. All in all, we report on redox-driven modulation of LINC-complexes integrity and on the identification of the first mammalian autophagy receptor which coordinately act to regulate dynamics of the mammalian NE. Our results launch molecular dissection of NE dynamics in health and disease.

## Acknowledgments

We thank the members of Molinari’s laboratory for discussions, and critical reading of the manuscript. We also thank Euro-BioImaging (www.eurobioimaging.eu) for providing access to imaging technologies and services via the Italian Node (ALEMBIC, Milan-Italy), Mihajlo Vanevic, Benjamin A. Barad, Miguel R. Leung and Gonzalo Obal for help with data processing and Andreas F. Sonnen for initial help with FIB milling. We are grateful to Stuart C. Howes and Menno Bergmeijer for cryo EM support as well as to Mariska Gröllers-Mulderij for cell culture support. M.M. is supported by the ALPHA-1 Foundation Research Grant (ID: 681136), the Foundation for Research on Neurodegenerative Diseases, the Swiss National Science Foundation (SNF, 310030_184827/2), the Eurostar (E! 113321 – CHAPERONE), Innosuisse (35449.1 IP-LS), and the Comel and Gelu Foundations. The work was also supported by the European Research Council under the European Union’s Horizon 2020 Program (ERC Consolidator Grant Agreement 724425 - BENDER) and the Nederlandse Organisatie voor Wetenschappelijke Onderzoek (Vici 724.016.001 to FF and Veni 212.152 to JF).

## Author contribution

M.K.K and J.F. performed the experiments, and the biochemical, imaging, transcriptional and quantitative analyses; C.G. and T.S. prepared DNA constructs and performed qPCR and biochemical analyses; D.M. assisted in CSLM analyses and set up and adapted the LysoQuant program. A.R. performed the analyses with the RT-TEM. J.F. and F.F. performed the FIB-CET analyses. M.M. designed the study, analyzed data, and wrote the manuscript. All the authors discussed the results and the manuscript.

The authors declare no competing interests.

## Correspondence and material requests

**Supplementary Fig. 1.**
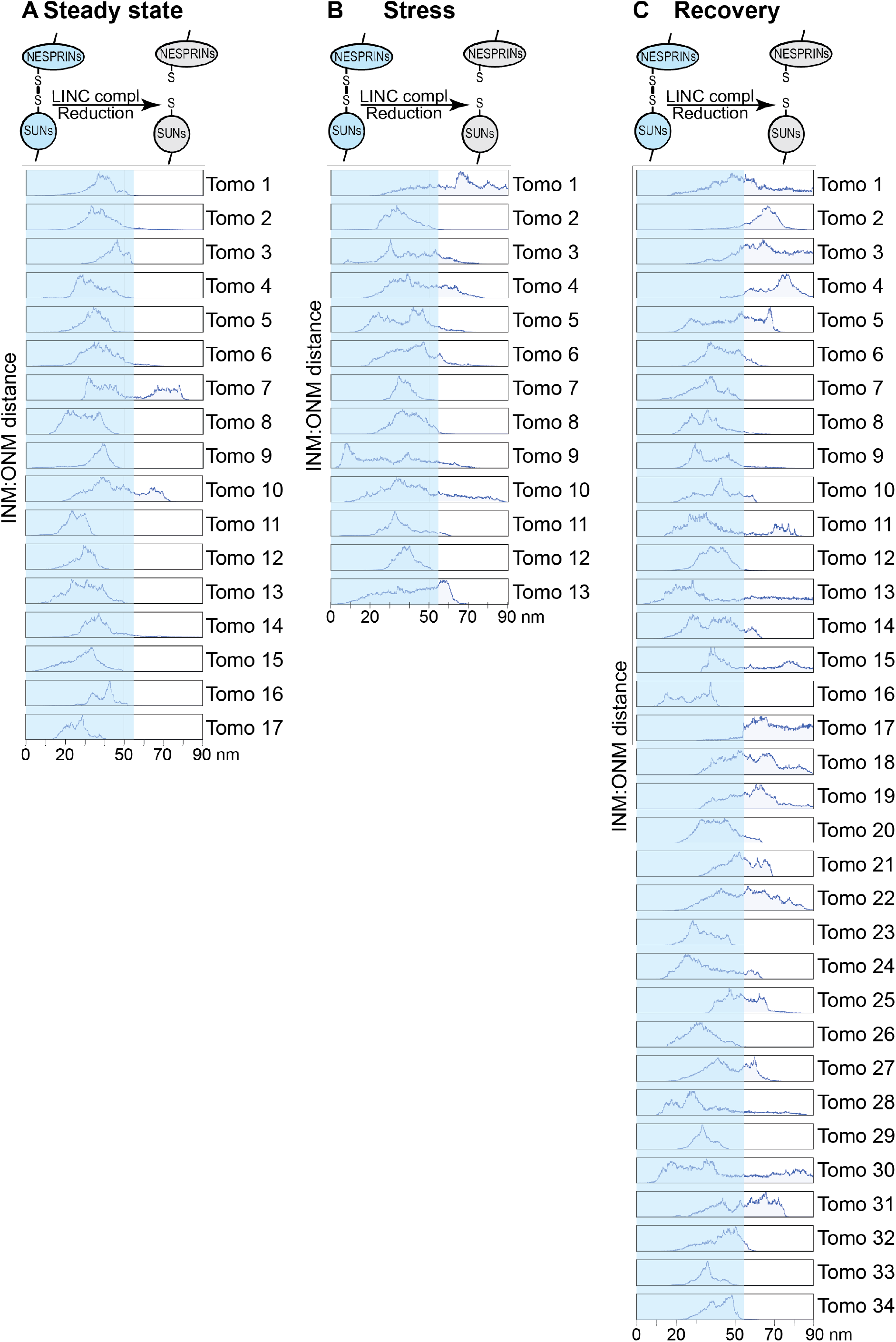
**A** The distance between INM and ONM was measured in 17 tomograms (steady state). **B** As **A** for 14 tomograms (pharmacologic induced ER stress). **C** Same as **A** for 34 tomograms (images of nuclei collected 5 h after interruption of the CPA treatment).

**Supplementary Fig. 2.**
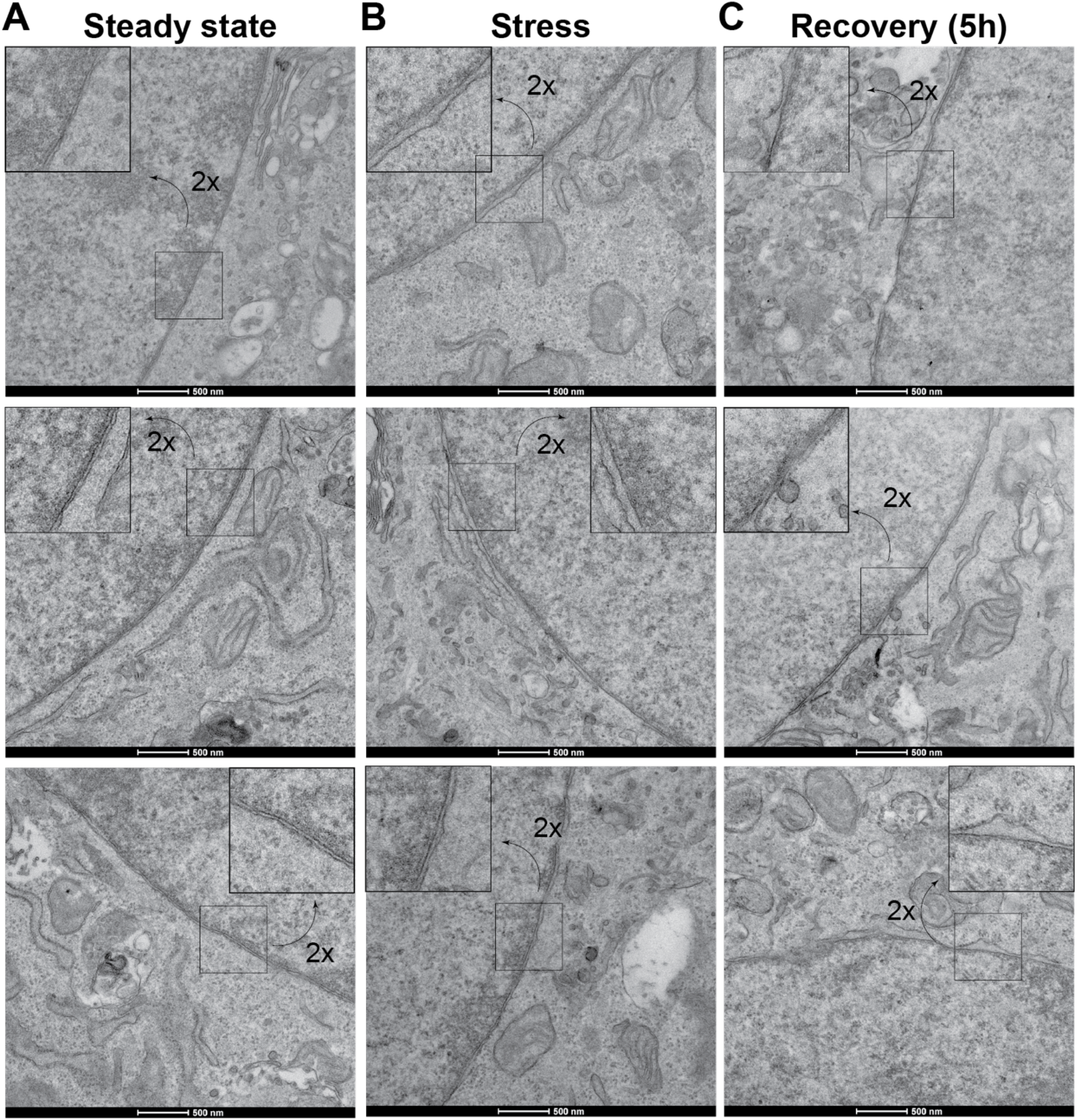
**A** Additional examples of NE ultrastructure at steady state, as in **Fig. 1H**. **B** Additional examples of NE ultrastructure during pharmacologic ER stress induction, as in **Fig. 1I**. **C** Additional examples of NE ultrastructure 5 h after interruption of the pharmacologic treatment, as in **Figs. 1J-1K**.

**Supplementary Fig. 3.**
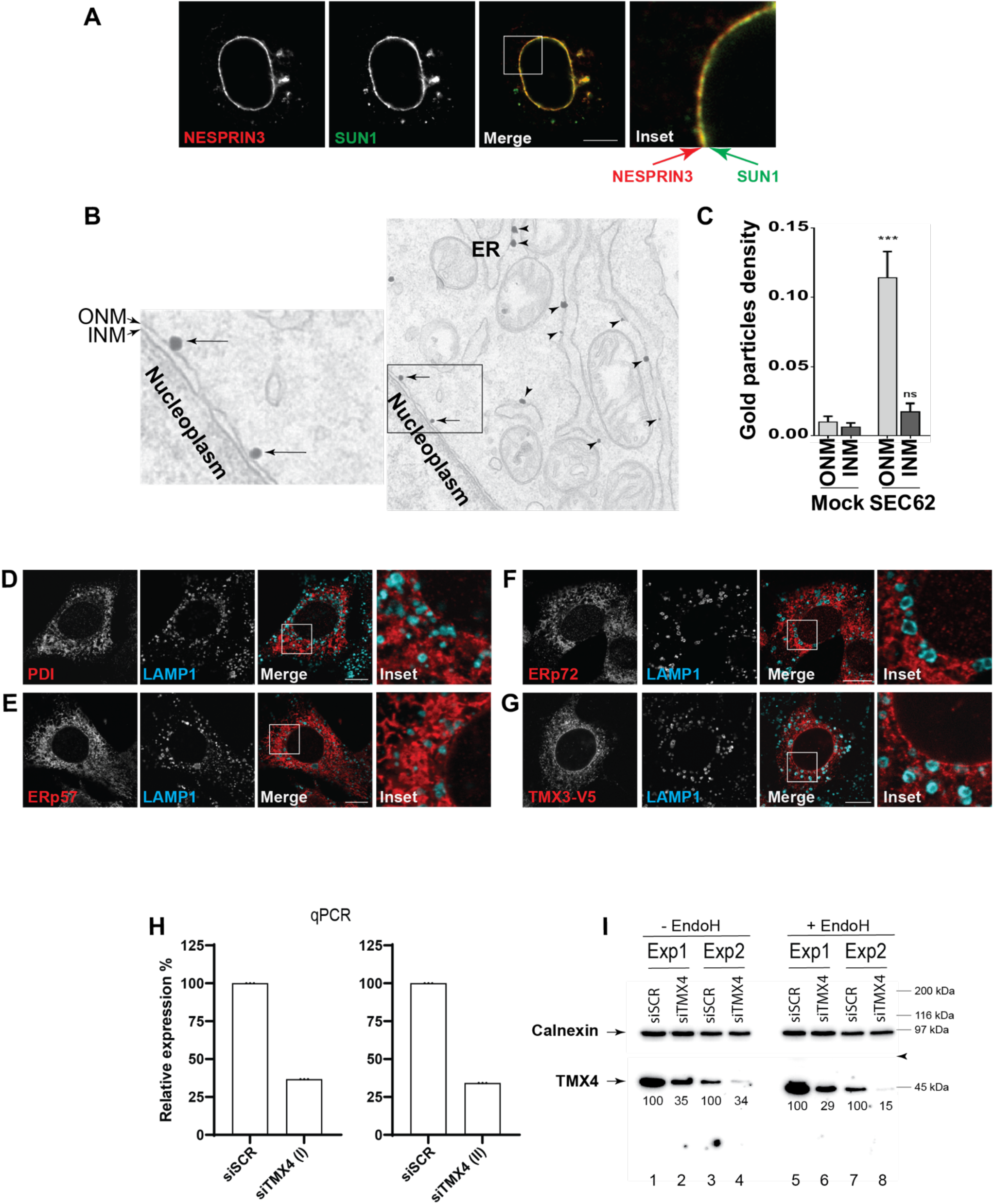
**A** Control showing the sub-cellular localization of the ONM marker protein NESPRIN3 and the INM marker protein SUN1 ^21^. **B** Micrograph showing immunogold labeling of endogenous SEC62 in the ER and in the ONM. **C** Distribution of gold particles in the ONM and in the INM of samples, where the primary antibody (anti SEC62) was not used (Mock) and in samples where the antigen is labeled with gold (SEC62). **D** Subcellular distribution of the PDI family member PDI in the ER. **E** Same as **D** for ERp57. **F** Same as **D** for ERp72. **G** Same as **D** for ectopically expressed TMX3-V5. **H** Control of the interference with TMX4 expression obtained by specific siRNas showing a 70% reduction of TMX4 transcript as determined with two different TMX4-specific primers. **I** Same as **H** to control reduction of TMX4 at the protein level in two separate experiments. The ER resident protein calnexin is shown as reference (lanes 1-4). The WB analysis has also been performed upon treatment of the cell lysate with endoglycanase H to remove the N-glycan that could interfere with antibody detection (lanes 5-8). Relative intensity of the TMX4 protein bands is given. Arrowhead shows where the membrane has been cut. The upper part has been probed with an anti-calnexin, the lower part with an anti-TMX4 antibody.

**Supplementary Fig. 4.**
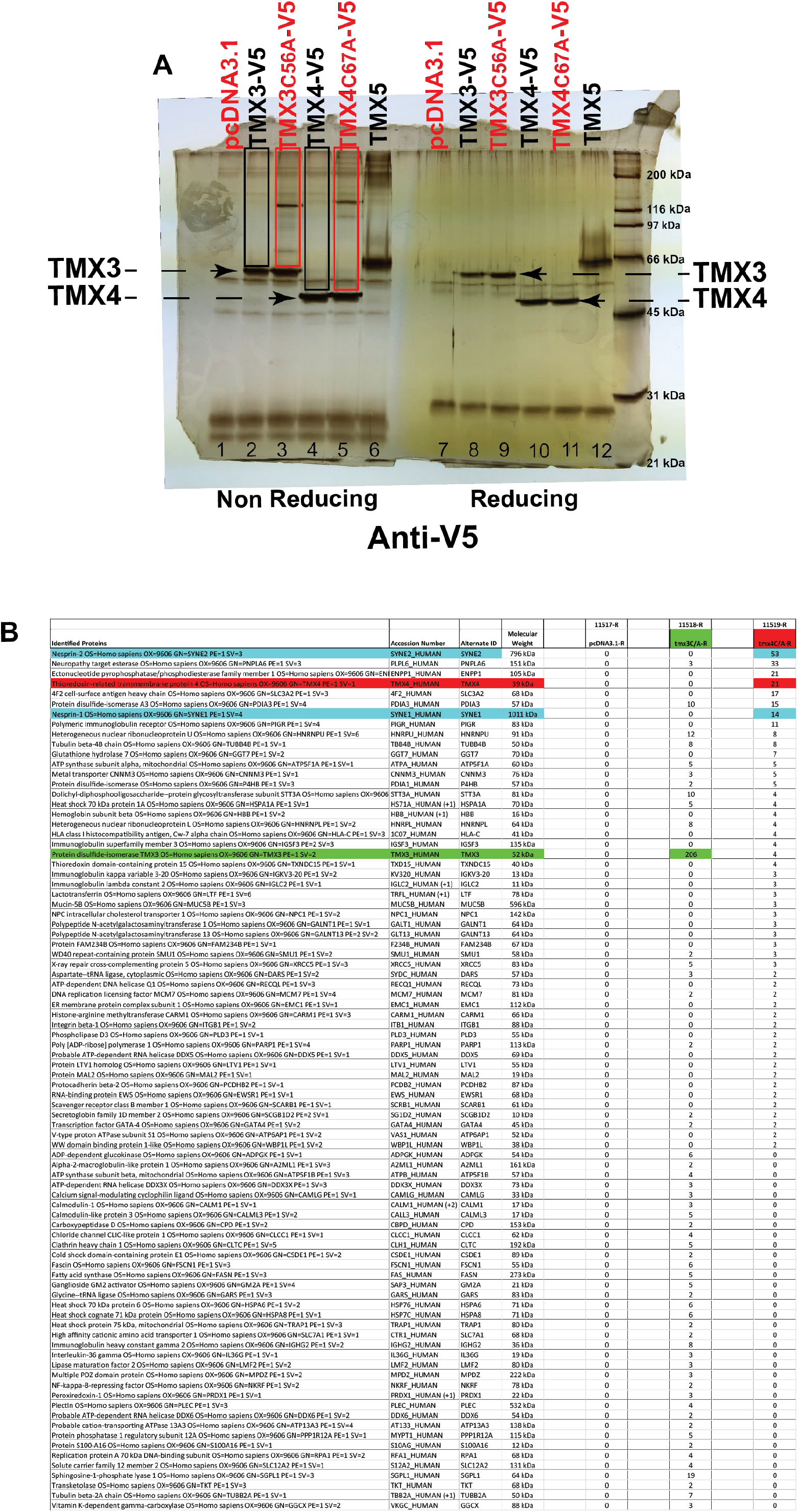
**A** Silver-stained gel showing TMX3 (lanes 2 and 8, as separated in non-reducing or reducing gels, respectively), TMX3_C56A_ (lanes 3 and 9), TMX4 (lanes 4 and 10) and TMX4_C67A_ (lanes 5 and 11). The polypeptide bands in the rectangle are mixed disulfides containing clients (i.e., cellular proteins covalently associated with the corresponding oxidoreductases that were identified by mass spectrometry). These complexes are disassembled, and are not visible anymore, upon sample reduction (lanes 7-12). Lanes 6 and 12 show TMX5 and its clients. **B** Cellular polypeptides engaged in mixed disulfides with TMX3 (green) or TMX4 (red) as identified in mass spectrometry. NESPRIN proteins are shown in blue in the table.

## Material and Methods

### Antibodies, Expression plasmids

Commercial antibodies used in this study: Lamp1 (DSHB Hybridoma Product 1D4B deposited by J. T. August, Immunofluorescence (IF) 1:50), V5 (Western blot (WB) 1:5000 Abcam), NESPRIN3 (WB 1:500 Abcam). Alexa–conjugated secondary antibodies were purchased from Thermo Fisher Scientific (IF 1:300). The HRP–conjugated secondary antibodies were purchased from SouthernBiotech (mouse, WB 1:20 000) and Protein A HRP–conjugated were purchased from Invitrogen (WB 1: 20 000). The HALO-Trap Agarose beads and V5-conjugated beads were purchased from ChromoTek and Sigma, respectively.

Antibody against SEC62 (IF 1:100) is a kind gifts from R. Zimmermann. Plasmid encoding GFP-RAB7, GFP-SUN1, GFP-NESPRIN3 are kind gifts from T. Johansen, H. J. Worman and A. Sonnenberg, respectively. NESPRIN3 was subcloned in a pcDNA3.1 expression plasmid with the addition of a N-terminal HaloTag. The trapping mutants of TMX4 (TMX4_C67A_-V5) and TMX3 (TMX3_C53A_-V5) were synthesized by GenScript. The TMX3-V5 WT and TMX4-V5 WT were obtained from trapping mutants by site-directed mutagenesis.

### Cell culture, transient transfection, CRISPR/Cas9 genome editing, small interfering RNA silencing

MEF, HEK293 and NIH/3T3 cells were grown in DMEM with 10% fetal bovine serum (FBS) at 37°C and 5% CO_2_. Transient transfections were carried out using the JetPrime transfection reagent (PolyPlus) in accordance with the manufacturer’s protocol. WT and *Atg5* KO MEF cells were kind gifts from M. Komatsu and N. Mizushima. CRISPRMOCK, CRISPRSEC62 and CRISPRFAM134B were generated by CRISPR/Cas9 genome editing in our lab as described in ^9^. RNA interferences were performed in MEF plated at 50-60% confluence. Cells were transfected with scrambled small interfering RNA or small interfering RNA (siRNA) to silence TMX4 expression (5′-GCAUGGUGUUCUUACGUUUtt-3′, 100 nM per dish, Silencer Select Pre-designed, Ambion). Cells were processed for immunofluorescence or for biochemical analyses after 48h of transfection.

### RNA extraction, polymerase chain reaction with reverse transcription (RT–PCR)

The extraction of RNA from MEF was performed with the GenElute Mammalian Total RNA Miniprep Kit (Sigma) according to the manufacturer’s instructions. One microgram of RNA was used for cDNA synthesis with dNTPs (Kapa Biosystems), oligo(dT) and the SuperScript II reverse transcriptase (Invitrogen) according to the instructions of the manufacturer. For each qRT-PCR reaction, 10 μl of Power SYBR Green PCR Master Mix (Bimake), 0.4 μl of reference ROX dye and 7.6 μl of milliQ sterile water were added to 1 μl cDNA together with 1 μl of 10 μM forward and reverse primer mix (mTMX4 (I) fwd: (5’-3’): TTG AGT GGC CGC TTC TTT GT rev: (5’-3’): CCA GAC ATC GTT AGA GAG GCT; mTMX4 (II) fwd: (5’-3’): CAT CCT GCC AGC AGA CTG ATT rev: (5’-3’): GGC GGA ATA TCC CAT CTT TTG C) for the transcript of interest in 96-well reaction plate (MicroAmp Fast Optical 96-Well Reaction Plate with Barcode (0.1 ml), Applied Biosystems). The plate was vortexed and centrifuged. Samples were loaded as triplicates. Quantitative real-time PCR was performed using QuantStudio™ 3 Real-Time PCR System. The housekeeping gene actin was used as reference. Data were analyzed using the QuantStudio™ Design & Analysis Software v1.5.5.

### Protocol to induce transient ER stress

To induce a transient ER stress, cultured cells were exposed for 12 h to 10 µM CPA in DMEM, 10% FBS. For the recovery condition, CPA was washed out and incubation was prolonged in DMEM, 10% FBS up to 48 h. Cells were processed for biochemical or imaging analyses (as in ^9,10^).

### Isolation of cell nuclei

HEK293 cells plated on poly-L-lysine coated dishes were washed in cold phosphate-buffered saline (PBS) containing 20 mM N-ethylmaleimide (NEM) and were collected by scraping in hypotonic lysis buffer (10 mM HEPES pH 7.9, 10 mM KCl, 1.5 mM MgCl_2_, 20 mM NEM and protease inhibitors cocktail (1 mM PMSF, chymostatin, leupeptin, antipain, and pepstatin, 10 μg/ml each)). NP-40 was added after 15 min of incubation on ice (0.5% final concentration). Cell lysates were vortexed gently, for 10 sec and incubated for 5min on ice. Nuclei were collected by centrifugation at 1500 rpm for 5min at 4°C. Pellet was washed once in hypotonic lysis buffer without PIs and centrifuged at 1500 rpm for 5min at 4 °C. Then, they were resuspended in RIPA buffer (1% Triton X-100, 0.1% SDS, 0.5% sodium deoxycholate in HBS, pH 7.4, 20 mM NEM and protease inhibitors), maintained for 20 min on ice and then homogenized with syringe every 5 min (BD Microlance™ 3 Needle 0,5 × 25 mm). Nuclear lysate supernatants were collected at 1000 rpm for 10 min at 4°C ^43^.

### Proteins immunoprecipitation and Western blot

The nuclear lysates were incubated with HALO-Trap Agarose beads for 2 h at 4°C to isolate HALO-NESPRIN3. After immunoprecipitation, samples were washed in 1 ml HBS 1x pH7.4, 0,5% Triton for 2 times. Beads were dried out and resuspended in sample buffer without DTT at 95°C for 5 min. Then, half of the sample was reduced by addition of 100 mM DTT. Samples were then boiled again 5 min at 95°C and loaded on 8% SDS-PAGE (polyacrylamide gel electrophoresis) for reducing and non-reducing protein separation. Proteins were transferred to polyvinylidene difluoride (PVDF) membranes with the Trans-Blot Turbo Transfer System (Bio-Rad). Membranes were blocked 10 min with 8% (w/v) non-fat dry milk (Bio-Rad) in Tris-buffered saline (TBS)-T and stained with primary antibodies (listed in material section) overnight, and for 45 min with HRP-conjugated secondary antibodies or Protein A HRP–conjugated. Membranes were developed using the Luminata Forte ELC detection system (Millipore) and signals were acquired with the ImageQuant LAS 4000 system (GE Healthcare Life Sciences). Membrane stripping was performed using Re-Blot Plus Strong Solution (Millipore) following the manufacturer’s instructions before repetition of the protocol for detection of other antigens.

### Silver staining

Polyacrylamide gels were fixed in a 40% EtOH, 10% Acetic Acid solution for 1-4 h at room temperature, rinsed two times for 20 min in 30% EtOH and then soaked for 20 min in H_2_0. After 1 min in 0.02% Na_2_S_2_O_3_ and 3 washes in H_2_0, the gels were incubated for 20 min in 0.2% AgNO_3_ and then rinsed 3 times in H_2_O. Development was performed with freshly prepared developing solution containing 3% Na_2_CO_3_ and 0.05% formaldehyde. Development was stopped by rinsing the gels in H_2_O and then in 5% Acetic Acid.

### Confocal laser scanning microscopy (CLSM)

MEF were seeded on alcian blue-treated glass coverslips and transiently transfected with JetPrime reagent according to the manufacturer’s protocol. Cells were then treated with 10 μM CPA for 12 h and, on CPA removal, incubated for 12 h with 50 nM BafA1 (Calbiochem), or DMSO (Sigma) and with 100 nM TMR HaloTag ligand (Promega). Following two PBS washes, cells were fixed with 3.7% formaldehyde diluted in PBS for 20 min at RT. Cells were permeabilized with 0.05% saponin, 10% goat serum, 10 mM HEPES, 15 mM glycine (PS) for 15 min. Cells were incubated with the primary antibodies diluted 1:100 in PS for 120 min, washed 2 times in PS, and then incubated with Alexa Fluor-conjugated secondary antibodies diluted 1:300 in PS for 45 min. Cells were rinsed 3 times with PS and once with water. Afterwards, cells were mounted with Vectashield (Vector Laboratories) with 4′,6-diamidino-2-phenylindole (DAPI).

Leica TCS SP5 microscope with a Leica HCX PL APO lambda blue 63.0 × 1.40 OIL UV objective was used to acquire confocal images.

The quantifications of NESPRIN3 or SUN1 positive lysosomes per cell were executed with LysoQuant, an unbiased and automated deep learning tool for fluorescent image quantification, which is freely available (https://www.irb.usi.ch/lysoquant/) ^22^.

### Live imaging

NIH3T3 cells were seeded on glass bottom Mattek 35 mm dishes (MatTek #1.5 coverslips) and transiently transfected with JetPrime reagent according to the manufacturer’s protocol for GFP-RAB7 and HALO-NESPRIN3. Cells were treated with 10 μM CPA for 12 h and, on CPA removal, incubated for 5 h with 100 nM BafA1, and with 100 nM Janelia Flour 646 HaloTag ligand (Promega).

Movies were recorded with a Leica TCS SP5 confocal microscope with a Leica HCX PL APO lambda blue 63.0 × 1.40 OIL UV objective was used to acquire confocal images at 37C, 5% CO2. Excitation was performed with 488 nm and 633 nm laser lines and fluorescence light was collected in ranges 493-625 nm and 637-791 nm respectively with pinhole 1 AU. The pictures were taken every 1 s with a pixel size of 63 nm and line average 3. Acquisitions were then cleaned with a median filter and movie processing was performed using Fiji/ImageJ.

### Mass spectrometry

HEK293 cells were transiently mock-transfected (pcDNA3.1) or transfected with TMX3-V5, TMX3_C53A_-V5, TMX4-V5, TMX4_C64A_-V5 and TMX5-V5. Sixteen h after transient transfection, cells were lysed and the PNS (post nuclear supernatant) was collected. The PNS was double immunoprecipitated using an anti-V5 conjugated beads. The beads used for immunoprecipitation of two independent experiments were dried out and processed for mass spectrometry analyses. Mass spectrometry analysis was performed at the Protein Analysis Facility, University of Lausanne, Lausanne, Switzerland.

### Statistical analyses and reproducibility

The legend of the figures indicates the number of independent experiments. Statistical analyses were performed using GraphPad Prism 9 software. Unpaired, two-tailed t test were used to asses statistical significance. P values are given in the figure legends; ns. P < 0.05; *P < 0.05; **P < 0.01; ***P < 0.001; ****P < 0.0001.

### Cryo electron tomography (CET): Grid preparation

MEF were seeded on R2/2 holey carbon on gold grids (Quantifoil or Protochips) coated with fibronectin in a glass bottom dish (Mattek or Ibidi) and incubated for 12 h. For the stress condition, cells were then incubated for 12 h with 10 µM CPA in DMEM + 10% FBS. For the recovery condition, CPA was washed out and cells were incubated in DMEM + 10% FBS for another 5 h. Grids were mounted to a manual plunger, blotted from the back for ∼10 s and plunged into liquid ethane.

### CET: Lamellae preparation

Lamellae were prepared using an Aquilos FIB-SEM system (Thermo Fisher Scientific). Grids were sputtered with an initial platinum coat (10 s) followed by a 10 s gas injection system (GIS) to add an extra protective layer of organometallic platinum. Samples were tilted to an angle of 15° to 22° and 11 µm wide lamellae were prepared. The milling process was performed with an ion beam of 30 kV energy in 3 steps: 1) 500 pA, gap 3 µm with expansion joints, 2) 300 pA, gap 1 µm, 3) 100 pA, gap 500 nm. Lamellae were finally polished at 30-50 pA with a gap of 200 nm.

### CET: Data acquisition

A total of 66 tilt series were acquired on a Talos Arctica (Thermo Fisher Scientific) operated at an acceleration voltage of 200 kV and equipped with a K2 summit direct electron detector and 20eV slit energy filter (Gatan). Images were recorded in movies of 5-8 frames at a target defocus of 4 to 6 μm and an object pixel size of 2.17 Å. Tilt series were acquired in SerialEM ^44^ using a grouped dose-symmetric tilt scheme ^45^ covering a range of ±60° with a pre tilt of ±10° and an angular increment of 3°. The cumulative dose of a series did not exceed 80 e-/Å^2^. On lower quality lamellae from CPA recovery condition, lower magnification tilt series were also acquired at an object pixel size of 4.47 Å, for membrane segmentation and illustration purposes (not used for distance measurements nor filament segmentation).

### CET: Tomogram reconstruction

Movie files of individual projection images were motion-corrected with MotionCor2 ^46^ and combined into stacks of tilt series using a Matlab script. The combined stacks were dose corrected in Matlab ^47^ and aligned using patch tracking in IMOD ^48^. Full tomograms were reconstructed by weighted back projection at a pixel size of 13.06 Å. Ice thickness was determined manually and was found to be <200 nm for all tomograms.

### CET: ONM to INM distance measurements

ONM and INM were manually segmented in Avizo software (Thermo Fisher Scientific) for each tomogram. The exported labels were submitted to surface morphometrics ^49^ to create a triangular mesh and measure the intra membrane distance along the surfaces. Results were plotted in Python using matplotlib.

### CET: Filament segmentation

The highest quality tomograms, corresponding to the thinner lamellae, were used for analysis of the filaments in the perinuclear region. Deconvolution was applied using tom_deconv in Matlab (https://github.com/dtegunov/tom_deconv) and the resulting tomograms were imported in Avizo software (Thermo Fisher Scientific). ONM and INM were manually segmented. Manual density-based segmentation was further used to trace the filaments visible in the perinuclear space in each z slice of the deconvoluted tomogram. The resulting labels were exported for visualization in UCSF Chimera ^50^.

### Room Temperature-Transmission Electron Microscopy (RT-TEM)

MEFs were seeded on alcian blue-coated glass coverslips and fixed in double strength fixative into the media (4% PFA EM grade, 5% GA in Na-cacodylate buffer 0.1 M, pH7.4) for 20 min at RT. After removing the mixture, cells were incubated with single strength fixative (2% PFA, 2.5% GA in Na-cacodylate buffer 0.1 M, pH7.4) for 3 h at RT. After several washes in cacodylate buffer, cells were post-fixed in 1% osmium tetroxide (OsO_4_), 1,5% potassium ferricyanide (K_4_[Fe(CN)_6_]) in 0,1 M Na-cacodylate buffer for 1 h on ice, washed with distilled water (dH_2_O) and enbloc stained with 0.5% uranyl acetate in dH_2_O overnight at 4°C in the dark. Samples were rinsed in dH_2_O, dehydrated with increasing concentrations of ethanol, embedded in Epon resin and cured in an oven at 60°C for 48 h. Ultrathin sections (70–90 nm) were collected using an ultramicrotome (UC7, Leica microsystem, Vienna, Austria), stained with uranyl acetate and Sato’s lead solutions and observed in a Transmission Electron Microscope Talos L120C (FEI, Thermo Fisher Scientific) operating at 120 kV. Images were acquired with a Ceta CCD camera (FEI, Thermo Fisher Scientific). Image analysis was performed using Microscopy Image Browser ^51^.

### RT-TEM: Immunogold electron microscopy

Cells were fixed in 4% PFA EM grade and 0.2 M HEPES buffer for 1 h at RT. After three washes in PBS, cells were incubated 10 min with 50 mM glycine and blocked 1 h in blocking buffer (0.2% bovine serum albumin, 5% goat serum, 50 mM NH_4_Cl, 0.1% saponin, 20 mM PO_4_ buffer, 150 mM NaCl). Staining with primary antibodies and nanogold-labeled secondary antibodies (Nanoprobes) were performed in blocking buffer at RT. Cells were fixed 30 min in 1% GA and nanogold was enlarged with gold enhancement solution (Nanoprobes) according to the manufacturer’s instructions. Cells were post fixed with OsO_4_ and processed as described for conventional EM. Images were acquired with Talos L120C TEM (FEI, Thermo Fisher Scientific) operating at 120 kV. Images were acquired with a Ceta CCD camera (FEI, Thermo Fisher Scientific).

### RT-Electron Tomography (RT-ET)

For electron tomography, 130-150 nm thick sections were collected on formvar-coated copper slot grids and gold fiducials (10 nm) were applied on both surfaces of the grids. The samples were imaged in a 120 kV Talos L120C TEM (FEI, Thermo Fisher Scientific). Single or dual tilted images series (+60/-60) were acquired using Tomography 4.0 software (FEI, Thermo Fisher Scientific). Tilted series alignment and tomography reconstruction were done with the IMOD software package ^52^. Segmentation and 3D visualization was done with Microscopy Image Browser ^51^ or IMOD software packages.

### RT-TEM: Measurement of INM-ONM distance

ONM and INM were manually segmented in Microscopy Image Browser (MIB) ^51^. Using the cell wall thickness plugin of MIB, the distance between the INM and the ONM was measured. Results were plotted in GraphaPad PRISM 9.

## Notes

### Competing Interest Statement

The authors have declared no competing interest.

